# Multiple timescales of sensory-evidence accumulation across the dorsal cortex

**DOI:** 10.1101/2020.12.28.424600

**Authors:** Lucas Pinto, David W. Tank, Carlos D. Brody

**Author notes:** corresponding authors: David W. Tank, Carlos D. Brody. Department of Physiology, Feinberg School of Medicine, Northwestern University, Chicago, IL, 60611 USA. these senior authors contributed equally to this work (David W. Tank, Carlos D. Brody).

## Abstract

Cortical areas seem to form a hierarchy of intrinsic timescales, but whether this is causal to cognitive behavior remains unknown. In particular, decisions requiring the gradual accrual of sensory evidence over time recruit widespread areas across this hierarchy. Here, we causally tested the hypothesis that this recruitment is related to the intrinsic integration timescales of these widespread areas. We trained mice to accumulate evidence over seconds while navigating in virtual reality, and optogenetically silenced the activity of many cortical areas during different brief trial epochs. We found that the inactivation of different areas primarily affected the evidence-accumulation computation per se, rather than other decision-related processes. Specifically, we observed selective changes in the weighting of evidence over time, such that frontal inactivations led to deficits on longer timescales than posterior cortical ones. Likewise, large-scale cortical Ca^2+^ activity during task performance displayed different temporal integration windows matching the effects of inactivation. Our findings suggest that distributed cortical areas accumulate evidence by leveraging their hierarchy of intrinsic timescales.

## Introduction

The cerebral cortex of both rodents and primates appears to be organized in a hierarchy of intrinsic integration timescales, whereby frontal areas integrate input over longer time windows than sensory areas (Cavanagh et al., 2020; Chaudhuri et al., 2015; Gao et al., 2020; Hasson et al., 2008; Ito et al., 2020; Kiebel et al., 2008; Murray et al., 2014; Runyan et al., 2017; Soltani et al., 2021; Spitmaan et al., 2020). Although this idea has received increasing attention, there is still no causal evidence that such timescale hierarchy is relevant for cognitive behavior.

In particular, the decisions we make in our daily lives often unfold over time as we deliberate between competing choices. This raises the possibility that decisions co-opt the cortical timescale hierarchy such that different cortical areas integrate decision-related information on distinct timescales. A commonly studied type of time-extended decision making happens under perceptual uncertainty, which requires the gradual accrual of sensory evidence (Bogacz et al., 2006; Brody and Hanks, 2016; Brunton et al., 2013; Carandini and Churchland, 2013; Gold and Shadlen, 2007; Morcos and Harvey, 2016; Newsome et al., 1989; Odoemene et al., 2018; Stine et al., 2020; Sun and Landy, 2016; Tsetsos et al., 2012; Waskom and Kiani, 2018). Neural correlates of decisions relying on evidence accumulation have been found in a number of cortical and subcortical structures, in both primates and rodents (Brincat et al., 2018; Ding and Gold, 2010; Erlich et al., 2015; Hanks et al., 2015; Horwitz and Newsome, 1999; Kim and Shadlen, 1999; Koay et al., 2020; Krueger et al., 2017; Murphy et al., 2020; Orsolic et al., 2021; Scott et al., 2017; Shadlen and Newsome, 2001; Wilming et al., 2020; Yartsev et al., 2018). Likewise, we have previously shown that, when mice must accumulate evidence over several seconds to make a navigational decision, the inactivation of widespread dorsal cortical areas leads to behavioral deficits, and that these areas encode multiple behavioral variables, including evidence (Pinto et al., 2019). However, we do not understand which aspects of these decisions lead to such widespread recruitment of brain structures.

Here, we hypothesized that the pattern of widespread recruitment of cortical areas during prolonged evidence accumulation can be explained by their underlying timescale hierarchy. To test this, we trained mice to accumulate evidence over seconds towards navigational decisions, and used brief optogenetic inactivation of single or combined cortical areas, restricted to one of six epochs of the behavioral trials. We show that the inactivation of widespread areas in the dorsal cortex affects primarily the evidence accrual process, rather than other decision-related computations. Further, the inactivation of different areas affects accumulation over distinct timescales, such that to an approximation frontal areas encode sensory evidence over longer temporal windows than posterior areas. In agreement with this, we show that cortical activity during the accumulation task displays a gradient of timescales, which are longer in frontal areas. Our findings thus suggest that evidence is accumulated by distributed cortical regions leveraging an existing hierarchy of temporal integration windows. Further, to our knowledge, they provide the first causal demonstration that this hierarchy is important for cognitive behavior.

## Results

### Brief inactivation of different cortical areas leads to accumulation deficits on distinct timescales

We trained mice to accumulate evidence over relatively long timescales while navigating in virtual reality (VR)(Figure 1A)(Pinto et al., 2018). The mice navigated a 3 m-long virtual T-maze and during the first 2 m (∼4 s) they encountered salient objects, or towers, along the walls on either side, and after a delay of 1 m (∼2 s) turned into the arm corresponding to the highest perceived tower count. The towers were visible for 200 ms, and appeared at different positions in each trial, obeying spatial Poisson processes of different underlying rates on the rewarded and non-rewarded side. Compatible with our previous reports (Koay et al., 2020; Pinto et al., 2018, 2019), task performance was modulated by the difference in tower counts between the right and left sides (Figure 1B, n = 20). Crucially, beyond allowing us to probe sensitivity to sensory evidence, the task design decorrelated the position of individual towers from the animals’ position in the maze across trials. This allowed us to build a logistic regression model that used the net sensory evidence (Δ towers, or #R – #L) from each of four equally-spaced bins from the cue region to predict the choice the mice made. In other words, we inferred the weight of sensory evidence from different positions in the maze on the final decision. While individual mice showed different evidence-weighting profiles, fitting the model on aggregate data yielded a flat evidence-weighting curve (Figure 1C, n = 100,787 trials), indicating that on average the mice weighted evidence equally from throughout the maze (Pinto et al., 2018). Note that all of the analyses presented below are performed on aggregate data (combined across mice, see Materials and Methods), such that our baseline condition is of even evidence weighting throughout the maze.

**Figure 1.**
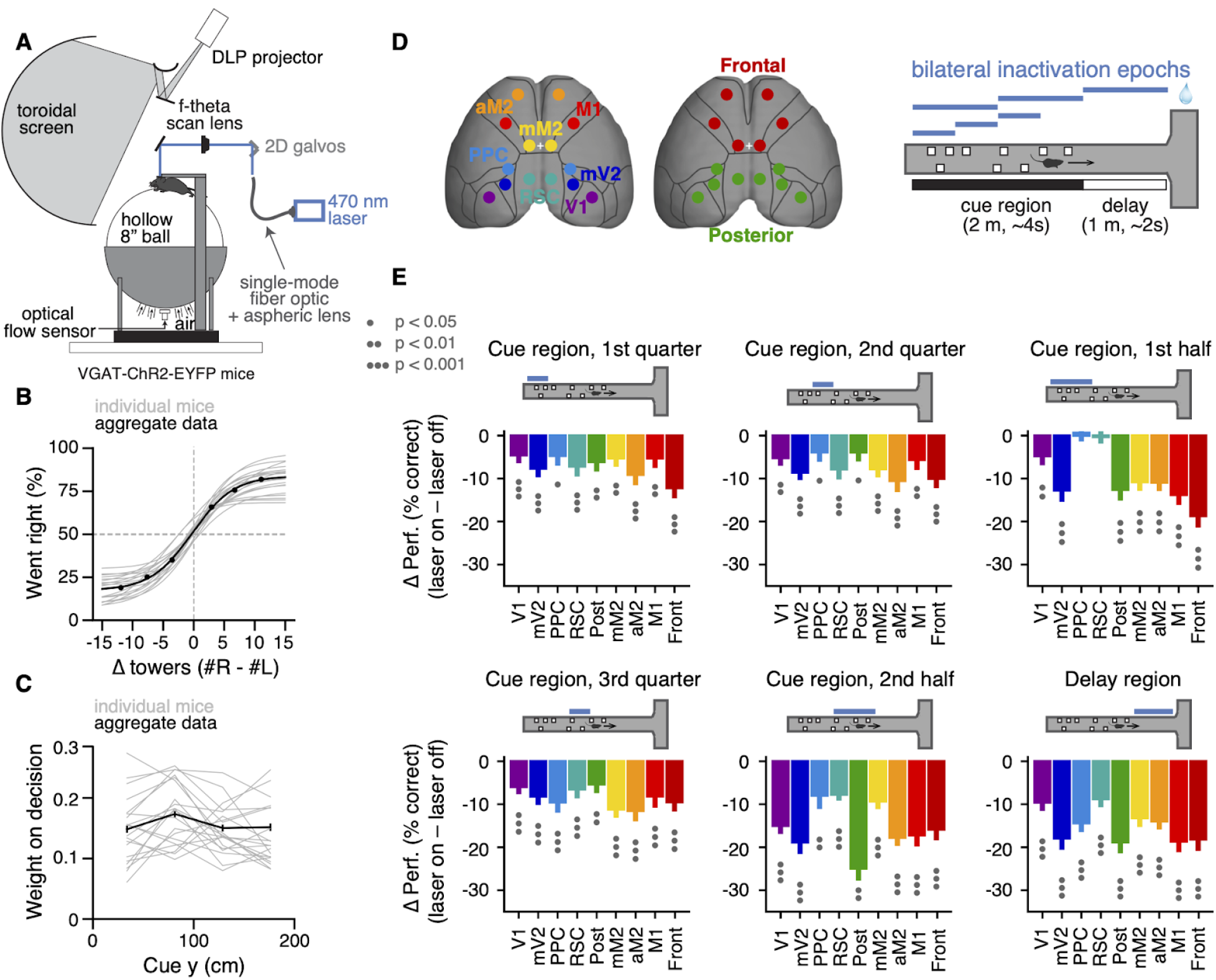
Temporally specific inactivation of multiple dorsal cortical regions during performance of a VR-based evidence-accumulation task. (**A**) Schematics of the experimental set-up. (**B**) Psychometric functions for control trials, showing the probability of right-side choice as a function of the strength of right sensory evidence, Δ towers (#R – #L). Thin gray lines: best-fitting psychometric functions for each individual mouse (n = 20). Black circles: aggregate data (n = 100,787 trials), black line: fit to aggregate data, error bars: binomial confidence intervals. (**C**) Logistic regression curves for the weight of sensory evidence from four equally-spaced bins on the final decision, from control trials. Thin gray lines: individual animals, thick black line: aggregate data, error bars: ± SD from 200 bootstrapping iterations. (**D**) Experimental design. We bilaterally inactivated 7 dorsal cortical areas, alone or in combination, while mice performed the accumulating-towers task. Bilateral inactivations happened during one of six regions in the maze spanning different parts of the cue region or delay. We thus tested a total of 54 area-epoch combinations. (**E**) Effects of subtrial inactivations on overall performance during all 54 area-epoch combinations. Each panel shows inactivation-induced change in overall % correct performance for each inactivation epoch, for data combined across mice. Error bars: S.D. across 10,000 bootstrapping iterations. Circles indicate significance according to the captions on the leftmost panel.

Our previous results have shown that cortical contributions to the performance of this task are widespread (Pinto et al., 2019), but our whole-trial inactivations did not allow us to tease apart the nature of the contributions of different areas. Here, we addressed this by asking how different dorsal cortical regions contribute to the weighting of sensory evidence in order to make a perceptual decision. To do this we cleared the intact skull of mice expressing Channelrhodopsin-2 (ChR2) in inhibitory interneurons (VGAT-ChR2-EYFP, n = 20), and used a scanning laser system to bilaterally silence different cortical regions, by activating inhibitory cells (Figure 1D) (Guo et al., 2014; Pinto et al., 2019). We targeted 7 different areas – primary visual cortex (V1), medial secondary visual cortex (mV2, roughly corresponding to area AM), posterior parietal cortex (PPC), retrosplenial cortex (RSC), the posteromedial portion of the premotor cortex (mM2), the anterior portion of the premotor cortex (aM2), and the primary motor cortex (M1) – as well as two combinations of these individual ares, namely posterior cortex (V1, mV2, PPC and RSC) and frontal cortex (mM2, aM2 and M1). Cortical silencing occured in one of six trial epochs: 1^st^, 2^nd^ or 3^rd^ quarter of the cue region (0 – 50 cm, 50 – 100 cm or 100 – 150 cm, respectively), 1^st^ or 2^nd^ half of the cue region (0 – 100 cm or 100 – 200 cm, respectively), or delay region (200 – 300 cm). We tested all the 54 possible area-epoch combinations (Figure 1–table supplement 1). This large number of experimental conditions allowed us to assess how the inactivation of different areas affects the use of current or past sensory evidence towards a final decision.

Compatible with our previous whole-trial inactivation experiments (Pinto et al., 2019), we found that the inactivation of all tested cortical areas significantly affected behavioral performance, though to varying degrees (Figure 1E, Figure 1–figure supplement 1). Furthermore, we observed a variety of effect profiles across regions and inactivation epochs, as assessed by the difference between the evidence-weighting curves separately calculated for ‘laser off’ and ‘laser on’ trials (Figure 1–figure supplement 2). Different effects were observed even comparing regions that were in close physical proximity (e.g. V1 and mV2). Additionally, all tested areas had significant effects in at least a subset of conditions (Figure 1–figure supplement 2, p<0.05, bootstrapping).

Most changes in the evidence-weighting curves happened for evidence concomitant to or preceding laser onset, indicating that the manipulations primarily affected the processing and/or memory of the evidence, i.e. the accumulation process itself (Figure 1–figure supplement 2). To quantify this, for each cortical area we aligned the control-subtracted evidence-weighting curves from different inactivation epochs by the position of laser onset, and focused on the changes in weights of evidence occurring up to 100 cm in the past (∼2s, Figure 2A). While some variability in these laser-onset-triggered curves suggests that the effects of inactivation depend somewhat on the exact inactivation epoch, the aligned curves from different epochs were fairly consistent (Figure 2B, gray lines), providing a concise summary of the multiple experimental conditions. Interestingly, the effects varied systematically according to the inactivated region. For example, mV2 inactivation led to a significant drop in the weight of evidence occurring while the laser was on (p = 0.004, one-sided paired t test), a trend towards affecting the memory of evidence occurring 50 cm in the past (∼1 s, p = 0.045, not significant after false discovery rate correction), and no discernible effect on evidence occurring 100 cm in the past (∼2 s, p = 0.41). Conversely, aM2 inactivation led to significant decreases in weighting all evidence between 100 cm in the past and the time of laser onset (p < 0.05, one-sided paired t test), with no differences in magnitude between position bins (F_2,10_ = 0.27, p = 0.77, mixed-model one-way ANOVA). This lack of modulation of effect size across y position bins was also true for the two other frontal areas, mM2 and M1 (Figure 2B, p > 0.88). Thus, subsets of cortical areas resembled each other in terms of the effects of their inactivation. Indeed, they could be optimally grouped into three clusters using spectral clustering (Figure 2C). Cluster 1 contained all frontal areas in our dataset: M1, mM2 and aM2. On average, this cluster resembled the effects described for aM2 above. In other words, the inactivation of frontal areas tended to equally and significantly affect weights for evidence occurring up to 100 cm in the past, suggesting that these areas accumulate evidence at fairly long timescales (p < 0.001 for all position bins, one-sided paired t test). Cluster 2 contained mV2 and PPC, and on average showed monotonically decreasing effects of inactivation on the weight of evidence as it gets more distal from laser onset (p < 0.02 for 0 and 50 cm, p = 0.47 for 100 cm). Thus, compared to the frontal area cluster, these posterior areas contributed to evidence accumulation on shorter timescales. Finally, cluster 3 contained V1 and RSC, whose inactivation led to non-monotonic changes in evidence weighting, affecting current and long-past evidence (p < 0.001), but not evidence occurring in between (p = 0.07). This is potentially compatible with findings that multiple timescales of processing can be present within the same cortical regions (Bernacchia et al., 2011; Cavanagh et al., 2020; Scott et al., 2017; Spitmaan et al., 2020; Wasmuht et al., 2018). Note that, for stimuli occurring while the laser is on, our analysis does not allow us to differentiate between pure visual processing deficits and deficits in evidence accumulation. However, this confound does not affect our main conclusions, since the areas also differ in terms of inactivation effects on evidence occurring prior to laser onset. Importantly, we have previously verified in an identical preparation that our laser parameters lead to robust inactivation and near-immediate recovery of pre-laser firing rates, with little to no rebound (Pinto et al., 2019). Thus, the effects observed here are unlikely to be related to changes in the average population activity levels outside of the nominal inactivation periods, or to different inactivation efficiencies between different epoch durations. Moreover, the effects were not due to increases in the timescale of the behavior itself leading to more forgetting, since we observed no significant laser-induced decreases in running speed (Figure 2–figure supplement 1).

**Figure 2.**
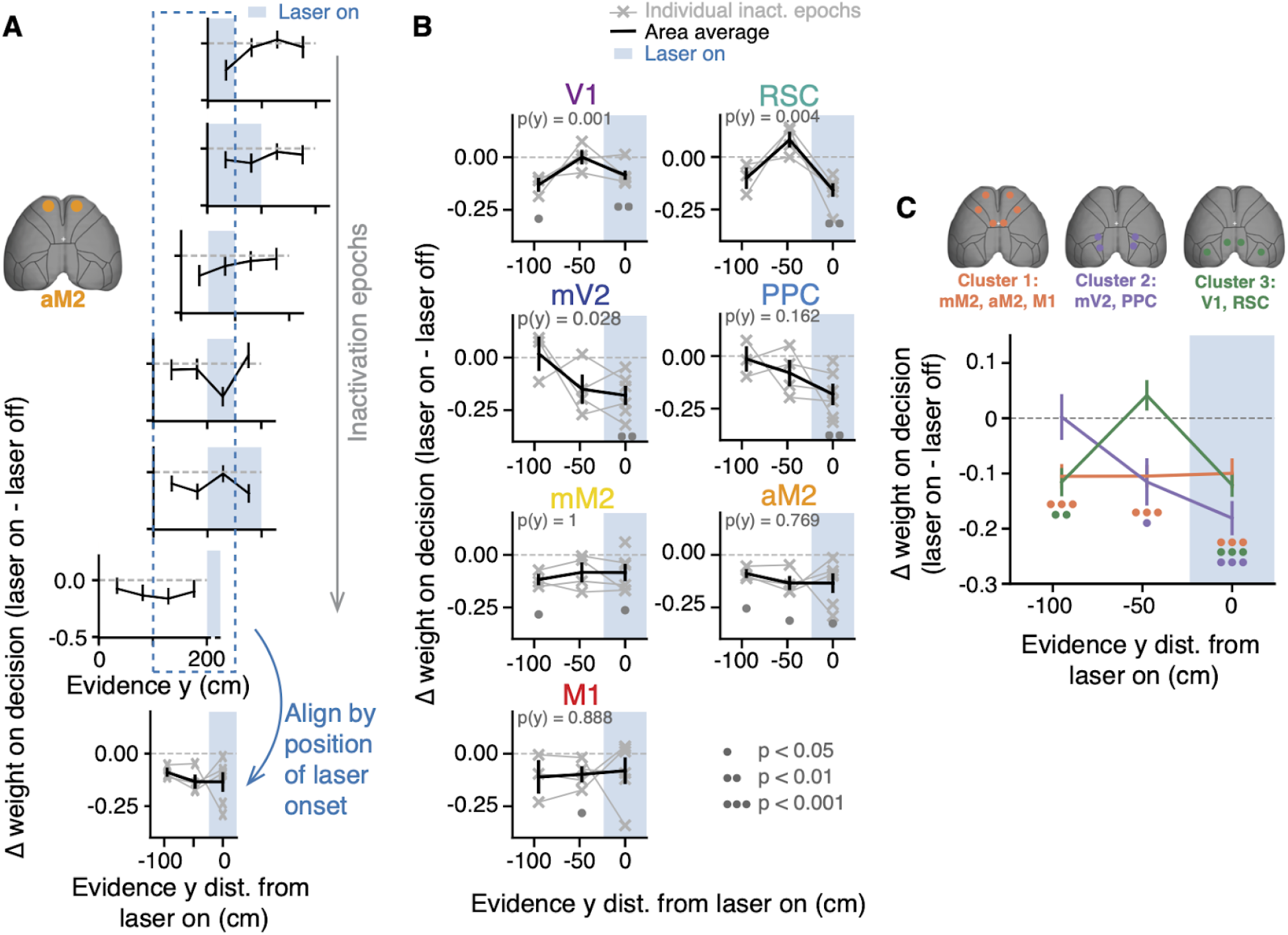
Inactivating different cortical areas leads to evidence-accumulation deficits on distinct timescales. (**A**) Illustration of the analysis method presented in panels B and C, using area aM2 as an example. Top six plots: effects of inactivating area aM2 during different epochs on evidence-weighting curves (laser off - laser on). Blue shading: laser-on epoch, error bars: ± SD across 10,000 bootstrapping iterations, data combined across mice. Bottom plot: Six top plots are aligned by laser onset (gray lines), and combined to include the first data point during ‘laser on’, and two preceding data points. See panel B for conventions. Error bars, ± SEM across experimental conditions. (**B**) Laser-onset-aligned changes in evidence-weighting curves for each bilaterally targeted area (laser on - laser off). Thin gray lines, individual inactivation epochs (n = 6 for y = 0, 4 for y = 50 and 3 for y = 100). Thick black lines, average across conditions. Error bars, ± SEM across experimental conditions. Shaded areas indicate laser on. Circles below the lines indicate statistical significance according to the caption on the bottom (one-sided paired t test vs. zero, corrected for multiple comparisons). P-values on top of each panel are from a one-way ANOVA with repeated measures with different y positions as factors. (**C**) Average laser-onset-aligned changes in evidence-weighting curves for each custer (caption on top). Error bars, ± SEM across inactivation epochs concatenated for each cluster. Circles below the lines indicate statistical significance for the cluster of corresponding color, according to the caption in panel a (one-sided paired t test vs. zero, corrected for multiple comparisons).

Next, we performed a similar analysis to assess changes in evidence weights after laser offset. Confirming our initial impression, the inactivation of most areas did not impact evidence weighting prospectively (Figure 2–figure supplement 2). Interestingly, however, mV2 and mM2 inactivation led to moderately but significantly decreased use of evidence occurring 100 cm in the future (p < 0.05, one-sided paired t test), perhaps suggesting an additional role for these areas also in post-accumulation decision processes (Hanks et al., 2015). Thus, our inactivation data suggest that the widespread cortical involvement in this task is largely related to the accumulation of sensory evidence, and that different cortical areas accumulate on distinct timescales. We next wondered whether evidence information from different areas is linearly combined, at least from a behavioral standpoint. To do this, we compared the effects of simultaneously inactivating all frontal or posterior areas to that expected by a linear combination of the effects of inactivating areas individually (i.e. their average). Neither posterior nor frontal areas significantly deviated from the linear prediction (Figure 2–figure supplement 3, p > 0.05, two-way ANOVA with repeated measures, factors y position and inactivation type). This suggests that signals from the different dorsal cortical areas could be combined by downstream regions in a near-linear fashion. Candidate regions include the medial prefrontal cortex, or subcortical structures such as the striatum and the cerebellum, which have been shown to be causally involved in evidence accumulation (Deverett et al., 2019; Yartsev et al., 2018). Other subcortical candidates are midbrain regions shown to have a high incidence of choice signals in a contrast discrimination task (Steinmetz et al., 2019). A caveat here is that the high variance in the data may have masked small non-linearities that could be revealed with larger sample sizes. The possibility of a downstream integration of cortical signals is agnostic to this limitation, however.

### A hierarchy of timescales in large-scale cortical activity during evidence accumulation

Our inactivation results are reminiscent of the findings that cortical areas display a hierarchy of intrinsic timescales, such that primary sensory areas tend to integrate over shorter time windows than frontal and other association areas (Chaudhuri et al., 2015; Hasson et al., 2008; Murray et al., 2014; Runyan et al., 2017). While these are thought to arise in part from intrinsic cellular and circuit properties such as channel and receptor expression, amount of recurrent connectivity and relative proportions of inhibitory interneuron subtypes (Chaudhuri et al., 2015; Duarte et al., 2017; Fulcher et al., 2019; Gao et al., 2020; Wang, 2020), they appear to be modulated by task demands (Gao et al., 2020; Ito et al., 2020). Thus, to confirm whether this timescale hierarchy exists in the mouse cortex during performance of the accumulating-towers task, we reanalyzed previously published data consisting of mesoscale widefield Ca^2+^ imaging of the dorsal cortex through the intact cleared skull of mice expressing the Ca^2+^ indicator GCaMP6f in excitatory neurons (Figure 3A, Emx1-Ai93 triple transgenics, n = 6, 25 sessions)(Pinto et al., 2019). To do this, we enhanced our previous linear encoding model (or GLM) of the average activity of anatomically defined regions of interest (ROIs)(Pinto et al., 2019) by including two sets of predictors in addition to task events. First, for each ROI we added the zero-lag activity of other simultaneously imaged ROIs as coupling predictors, similar to previous work (Pillow et al., 2008; Runyan et al., 2017)(Figure 3–figure supplement 1). Crucially, we also included auto-regressive predictors to capture intrinsic activity auto-correlations that are unrelated to behavioral events. In other words, this approach allowed us to estimate within-task auto-correlations while separately accounting for task-induced temporal structure in cortical dynamics (Spitmaan et al., 2020). Adding these new sets of predictors resulted in a large and significant increase in cross-validated model accuracy, as measured by the linear correlation coefficient between the model predictions and a test dataset not used to fit the model (Figure 3B, C; ∼0.95 vs. ∼0.3, F_model(6,2,12)_ = 1994.85, p = 6.2 × 10^−13^, two-way ANOVA with repeated measures). Note that these values are computed on held-out raw data points rather than averaged activity. Thus, while the original model in our previous work had low cross-validated accuracies in comparison, those values are compatible with other encoding models of cortical activity in the literature that used similarly stringent goodness-of-fit metrics (e.g. Huth et al., 2012; Pinto and Dan, 2015).

**Figure 3.**
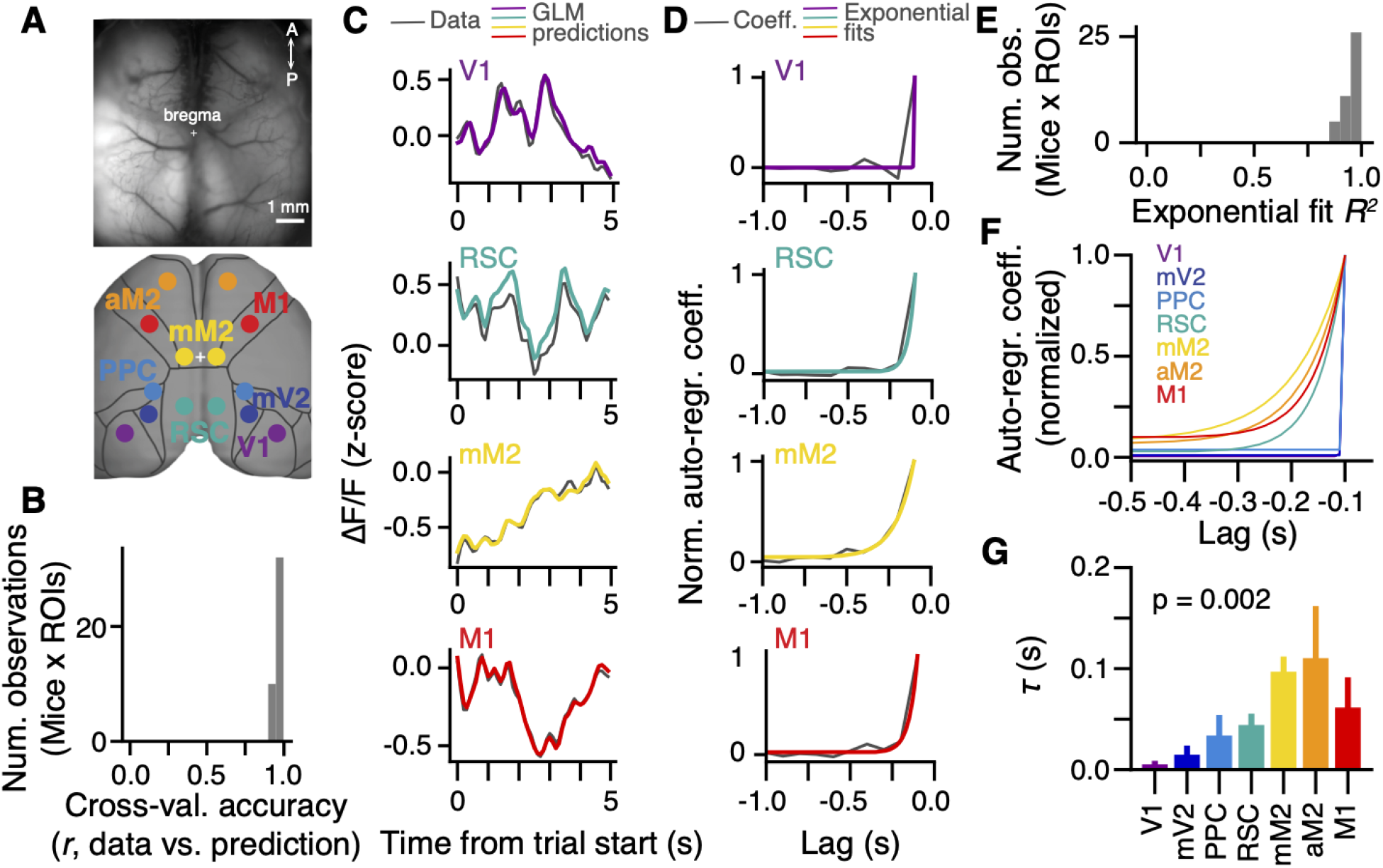
A hierarchy of activity timescales during evidence accumulation. (**A**) Top: example widefield imaging field of view showing GCaMP6f fluorescence across the dorsal cortex. Bottom: approximate correspondence between the field of view and ROIs defined from the Allen Brain Atlas, ccv3. (**B**) Distribution of cross-validated accuracies across mice (n = 6, sessions for each mouse are averaged) and ROIs (n = 7, averaged across hemispheres). (**C**) Example of actual ΔF/F (gray) and GLM predictions (colored lines) for the first 5 s of the same held-out single trial, and four simultaneously imaged ROIs. Traces are convolved with a 1-SD gaussian kernel for display only. (**D**) Auto-regressive GLM coefficients as a function of time lags for an example imaging session and four example ROIs. Gray, coefficient values. Colored lines, best-fitting exponential decay functions. (**E**) Distribution of *R2* values for the exponential fits across mice (n = 6, sessions for each mouse are averaged) and ROIs (n = 7, averaged across hemispheres). (**F**) Exponential decay functions for all seven cortical areas, fitted to the average across mice (n = 6). (**G**) Time constants extracted from the exponential decay fits, for each area. Error bars, ± SEM across mice (n = 6). P-value is from a one-way ANOVA with repeated measures with ROIs as factors.

Motivated by our inactivation findings, we focused our analysis on the auto-regressive coefficients of the model. We observed that across animals the rate of decay of these coefficients over lags slowed systematically from visual to premotor areas, with intermediate values for M1, PPC and RSC (Figure 3D). To quantify this, we fitted exponential decay functions to the auto-regressive coefficients averaged across hemispheres (Figure 3D–F), and extracted decay time constants (τ, Figure 3G). Compatible with our observations, τ differed significantly across cortical areas (F_6,30_ = 4.49, p = 0.002, one-way ANOVA with repeated measures), being larger for frontal than posterior areas, in particular PPC and mV2. Note that, while it is possible that these coefficients capture auto-correlations introduced by intrinsic GCaMP6f dynamics, there is no reason to believe that this affects our conclusions, as indicator dynamics should be similar across regions. Thus, during the evidence-accumulation task, cortical regions display increasing intrinsic timescales going from visual to frontal areas. This is consistent with previous reports for spontaneous activity and other behavioral tasks (Chaudhuri et al., 2015; Hasson et al., 2008; Murray et al., 2014; Runyan et al., 2017). Moreover, it is in overall agreement with our inactivation findings (Figure 2), and suggests that the different intrinsic timescales across the cortex support evidence integration over time windows of different durations.

## Discussion

Taken together, our results suggest that distributed cortical areas contribute to sensory-evidence accrual on different timescales. Specifically, brief sub-trial inactivations during performance of a decision-making task requiring seconds-long evidence accumulation resulted in distinct deficits in the weighting of sensory evidence from different points in the stimulus stream. This was such that, on average, the inactivation of frontal cortical areas resulted in decreased use of evidence occurring further in the past from laser onset compared to a subset of posterior regions (Figure 2). Compatible with this, using an encoding model of large-scale cortical dynamics, we found that activity timescales vary systematically across the cortex in a way that mirrors the inactivation results (Figure 3).

Our results add to a growing body of literature that has revealed that the cortex of rodents and primates appears to be organized in a hierarchy of temporal processing windows across regions (Chaudhuri et al., 2015; Gao et al., 2020; Hasson et al., 2008; Ito et al., 2020; Murray et al., 2014; Runyan et al., 2017; Spitmaan et al., 2020). Specifically, to the best of our knowledge, they provide the first causal demonstration that the contributions of different cortical areas to decision-making computations appear similarly arranged in a temporal hierarchy. A caveat here is that our inactivation findings do not exactly match the smooth increases in integration windows going from posterior to frontal areas that we observed in our neural data. Rather, they appear to reflect a more modular organization, as suggested by our clustering results (see also Pinto et al., 2019), and one that does not exactly map onto the expected monotonic effects on accrual timescales. The latter could be due the fact that diverse timescales exist at the level of individual neurons within each region (Bernacchia et al., 2011; Cavanagh et al., 2020; Scott et al., 2017; Spitmaan et al., 2020; Wasmuht et al., 2018), and/or that, other decision-making processes beyond evidence accumulation are also affected by our inactivations. For instance, both mV2 and mM2 appeared to contribute to post-accrual decision processes (Figure 2–figure supplement 2). Nevertheless, our results point to accrual timescale hierarchies being a significant factor explaining the large-scale functional organization of cortical dynamics during evidence-based decisions.

Our findings further suggest the possibility that the logic of widespread recruitment of cortical regions in complex, time-extended decisions may in part rely on intrinsic temporal integration properties of local cortical circuits, rather than specific evidence-accumulation mechanisms. For instance, it is possible that simple perceptual decisions primarily engage only the relevant sensory areas because they can be made on the fast intrinsic timescales displayed by these regions (Zatka-Haas et al., 2020). Along the same lines, it is conceivable that discrepancies in the literature regarding the effects of perturbing different cortical areas during evidence accumulation stem in part from differences in the timescales of the various tasks (Erlich et al., 2015; Fetsch et al., 2018; Hanks et al., 2015; Katz et al., 2016; Pinto et al., 2019).

An important remaining question is whether evidence from the different time windows is accumulated in parallel or as a feedforward computation going from areas with short to those with long integration time constants. The parallel scheme would be compatible with recent psychophysical findings in humans reporting confidence of their evidence-based decisions (Ganupuru et al., 2019). Conversely, a feedforward transformation would be in agreement with human fMRI findings during language processing (Yeshurun et al., 2017), and with a previously published model whereby successive (feedforward) convolution operations lead to progressively longer-lasting responses to sensory evidence (Scott et al., 2017). Interestingly, the oculomotor integrator of both fish and monkeys appears to be organized as largely feedforward chains of integration leading to systematically increasing time constants (Joshua and Lisberger, 2015; Miri et al., 2011), perhaps suggesting that this architecture is universal to neural integrators.

Much work remains before obtaining a complete circuit understanding of gradually evolving decisions. Our findings highlight the fact that, much like in memory systems (Jeneson and Squire, 2012), the timescale of decision processes is an important feature governing their underlying neural mechanisms, a notion which should be incorporated into both experimental and theoretical accounts of decision making.

## Materials and Methods

### Animals and surgery

All procedures were approved by the Institutional Animal Care and Use were performed in accordance with the Guide for the Committee at Princeton University and Care and Use of Laboratory Animals (National Research Council, 2011). We used both male and female VGAT-ChR2-EYFP mice aged 2 – 16 months [B6.Cg-Tg(Slc32a1-COP4*H134R/EYFP)8Gfng/J, Jackson Laboratories, stock # 014548, n = 28]. Part of the inactivation data from some of these animals was collected in the context of previous work (Pinto et al., 2019), but the analyses reported here are completely novel. The mice underwent sterile surgery to implant a custom titanium headplate and optically clear their intact skulls, following a procedure described in detail elsewhere (Pinto et al., 2019). Briefly, after exposing the skull and removing the periosteum, successive layers of cyanoacrylate glue (krazy glue, Elmers, Columbus, OH) and diluted clear metabond (Parkell, Brentwood, NY) were applied evenly to the dorsal surface of the skull, and polished after curing using a dental polishing kit (Pearson dental, Sylmar, CA). The headplate was attached to the cleared skull using metabond, and a layer of transparent nail polish (Electron Microscopy Sciences, Hatfield, PA) was applied and allowed to cure for 10 – 15 min. The procedure was done under isoflurane anesthesia (2.5% for induction, 1.5% for maintenance). The animals received two doses of meloxicam for analgesia (1 mg/kg I.P or S.C.), given at the time of surgery and 24 h later, as well as peri-operative I.P. injections of body-temperature saline to maintain hydration. Body temperature was maintained constant using a homeothermic control system (Harvard Apparatus, Holliston, MA). The mice were allowed to recover for at least 5 days before starting behavioral training. After recovery they were restricted to 1 – 2 mL of water per day and extensively handled for another 5 days, or until they no longer showed signs of stress. We started behavioral training after their weights were stable and they accepted handling. During training, the full allotted fluid volume was typically delivered within the behavioral session, but supplemented if necessary. The mice were weighed and monitored daily for signs of dehydration. If these were present or their body mass fell below 80% of the initial value, they received supplemental water until recovering. They were group housed throughout the experiment, and had daily access to an enriched environment (Pinto et al., 2018). The animals were trained 5 – 7 days/week.

The analysis reported in Figure 3 (widefield Ca^2+^ imaging) is from data collected in the context of a previous study (Pinto et al., 2019), although the analysis is novel. The data was from 6 male and female mice from triple transgenic crosses expressing GCaMP6f under the CaMKIIα promoter from the following two lines: Ai93-D;CaMKIIα-tTA [IgS6^tm93.1(tetO-GCaMP6f)Hze^ Tg(Camk2a-tTA)1Mmay/J, Jackson Laboratories, stock # 024108] and Emx1-IRES-Cre [B6.129S2-Emx1^tm1(cre)Krj^/J, Jackson Laboratories, stock # 005628]. These animals also underwent the surgical procedure described above.

### Virtual reality apparatus

The mice were trained in a virtual reality (VR) environment (Figure 1A) described in detail elsewhere (Pinto et al., 2018). Briefly, they sat on an 8-inch hollow Styrofoam® ball that was suspended by compressed air at ∼60 p.s.i, after passing through a laminar flow nozzle to reduce noise (600.326.5K.BC, Lechler, St. Charles, IL). They were head-fixed such that their snouts were aligned to the ball equator and at a height such that they could run comfortable without hunching, while still being able to touch the ball with their full paw pads (corresponding to a headplate-to-bal height of ∼1 inch for a 25-g animal). Ball movements were measured using optical flow sensors (ADNS-3080 APM2.6) and transformed into virtual world displacements using custom code running on Arduino Due (https://github.com/sakoay/AccumTowersTools/tree/master/OpticalSensorPackage). The ball sat on a custom 3D-printed cup that contained both the air outlet and the movement sensor. The VR environment was projected onto a custom-built toroidal Styrofoam® screen using a DLP projector (Optoma HD141X, Fremont, CA) at a refresh rate of 120 Hz and a pixel resolution of 1024 x 768. The screen spanned ∼270° of azimuth and ∼80° of elevation in the mouse’s visual field. The whole set-up was enclosed in a custom-built sound-attenuating chamber. The VR environment was programmed and controlled using ViRMEn (Aronov and Tank, 2014) (https://pni.princeton.edu/pni-software-tools/virmen), running on Matlab (Mathworks, Natick, MA) on a PC.

### Behavioral task

We trained the mice in the accumulating-towers task (Pinto et al., 2018). The mice ran down a virtual T-maze that was 3.3 m in length (y), 5 cm in height and a nominal 10 cm in width (x, though they were restricted to the central 1 cm). The length of the maze consisted of a 30-cm start region to which they were teleported at the start of each trial, followed by a 200-cm cue region and a 100-cm delay region. The cue and the delay region had the same wallpaper designed to provide optical flow. During the cue region, the mice encountered tall white objects (2 x 6 cm, width x height), or towers, that appeared at random locations in each trial at a Poisson rate of 7.7 m^-1^ and 2.3 m^-1^ on the rewarded on non-rewarded side, respectively (or 8.0 and 1.6 m^-1^ in some sessions), with a 12-cm refractory period and an overall density of 5 m^-1^. The towers appeared when the mice were 10 cm away from their drawn locations, and disappeared 200 ms later (roughly corresponding to the time over which the tower sweeps across the visual field given average running speeds). After the maze stem the mice turned into one of the two arms (10.5 x 11 x 5 cm, length x width x height), and received a reward if they turned to the arm corresponding to the highest tower count (4–8 µL of 10% v/v sweet condensed milk). This was followed by a 3-s inter-trial interval, consisting of 1 s of a frozen frame of the VR environment and 2 s of a black screen. An erroneous turn resulted in a loud sound and a 12-s timeout.

Each daily behavioral session (∼1 h, ∼200 – 250 trials) started with warm-up trials of a visually guided task in the same maze, in which towers appeared only on the rewarded side and additionally a 30-cm tall visual guide visible from the start of the trial was placed in the arm corresponding to the reward location. The animals progressed to the main task when they achieved at least 85% correct trials over a running window of 10 trials in the warm-up task. During the accumulating-towers task, performance was evaluated over a 40-trial running window, both to assess side biases and correct them using an algorithm described elsewhere (Pinto et al., 2018), and to trigger a transition into a 10-trial block of easy trials if performance fell below 55% correct. These blocks consisted of towers only on the rewarded side, and were introduced to increase motivation but were not included in the analyses. No optogenetic inactivation was performed during either warm-up or easy-block trials. In the widefield imaging experiments, the behavioral sessions contained several visually guided (warm up) blocks (Pinto et al., 2019). These were excluded from the present analyses.

### Laser-scanning optogenetic inactivation

We used a scanning laser setup described in detail elsewhere (Pinto et al., 2019). Briefly, a 473-nm laser beam (OBIS, Coherent, Santa Clara, CA) was directed to 2-D galvanometers using a 125-µm single-mode optic fiber optic (Thorlabs, Newton, NJ) and reached the cortical surface after passing through an f-theta scanning lens (LINOS, Waltham, MA). We used a 40-Hz square wave with an 80% duty cycle and a power of 6 mW measured at the level of the skull. This corresponds to an inactivation spread of ∼ 1.5 – 2 mm (Pinto et al., 2019). While this may introduce confounds regarding ascribing exact functions to specific cortical areas, we have previously shown that the effects of whole-trial inactivations at much lower powers (corresponding to smaller spatial spreads) are consistent with those obtained at 6 mW. To minimize post-inactivation rebounds, the last 100 ms of the laser pulse consisted of a linear ramp-down of power (Guo et al., 2014; Pinto et al., 2019). We performed inactivations during the following trial epochs: 1^st^, 2^nd^ or 3^rd^ quarter of the cue region (0 – 50 cm, 50 – 100 cm or 100 – 150 cm, respectively), 1^st^ or 2^nd^ half of the cue region (0 – 100 cm or 100 – 200 cm, respectively), or delay region (200 – 300 cm). Thus, the epochs were defined according to the animals’ y position in the maze. Because of this, the onset time of the power ramp-down was calculated in each trial based on the current speed and the expected time at which the mouse would reach the laser offset location. The system was controlled using custom-written code in Matlab running on a PC, which sent command analog voltages to the laser and galvanometers through NI DAQ cards. This PC received instructions for laser onset, offset and galvanometer position from the ViRMEn PC through digital lines.

We targeted a total of 9 area combinations, either consisting of homotopic bilateral pairs or multiple bilateral locations. The galvanometers alternated between locations at 200 Hz (20-mm travel time: ∼250 µs) and, in the case of more than 2 locations, the sequence of visited locations was chosen to minimize travel distance. The inactivated locations were defined based on stereotaxic coordinates using bregma as reference, as follows:

- Primary visual cortex (V1): –3.5 AP, 3 ML
- Medial secondary visual cortex (mV2, ∼ area AM): –2.5 AP, 2.5 ML
- Posterior parietal cortex (PPC): –2 AP, 1.75 ML
- Retrosplenial cortex (RSC): –2.5 AP, 0.5 ML
- Posteromedial portion of the premotor cortex (mM2): 0.0 AP, 0.5 ML
- Anterior portion of the premotor cortex (aM2): +3 AP, 1 ML
- Primary motor cortex (M1): +1 AP, 2 ML
- Posterior cortex: V1, mV2, PPC and RSC
- Frontal cortex: mM2, aM2 and M1

To ensure consistency in bregma location across behavioral sessions, the experimenter set bregma on a reference image and for each session the current image of the mouse’s skull was registered to this reference using rigid transformations. Different sessions contained different combinations of areas and inactivation epochs, resulting in partially overlapping mice and sessions for each condition. The probability of inactivation trials therefore varied across sessions, ranging from a total of 0.15 – 0.35 across conditions, and from 0.02 – 0.15 per condition. In our experience, capping the probability at ∼0.35 is important to maintain motivation throughout the behavioral session.

### Widefield Ca^2+^ imaging

Details on the experimental setup and data preprocessing can be found elsewhere (Pinto et al., 2019). Briefly, we used a tandem-lens macroscope (1x – 0.63x planapo, Leica M series, Wetzlar, Germany) with alternating 410-nm and 470-nm LED epifluorescence illumination for isosbestic hemodynamic correction, and collected 525-nm emission at 20 Hz, using an sCMOS (OrcaFlash4.0, Hamamatsu, Hamamatsu City, Japan), with an image size of 512 x 512 pixels (pixel size of ∼17 µm). Images were acquired with HCImage (Hamamatsu) running on a PC, and synchronized to the behavior using a data acquisition-triggering TTL pulse from another PC running ViRMEn, which in turn received analog frame exposure voltage traces acquired through a DAQ card (National Instruments, Austin, TX) and saved in the behavioral log file. The image stacks were motion-corrected by applying the x-y shift that maximized the correlation between successive frames, and then were spatially binned to a 128 x 128 pixel image (∼68 x 68 µm). The fluorescence values from pixels belonging to different anatomical ROIs were averaged into a single trace, separately for 410-nm (*F*_*v*_) and 470-nm excitation (*F*_*b*_). After applying a heuristic correction to *F*_*v*_ (Pinto et al., 2019), we calculated fractional fluorescence changes as *R* = *F/F*_*0*_, where *F*_*0*_ for each excitation wavelength was calculated as the mode of all *F* values over a 30-s sliding window with single-frame steps. The final ΔF/F was calculated using a divisive correction, *ΔF/F* = *R*_*b*_ / *R*_*v*_ – 1. ROIs were defined based on the Allen Brain Mouse Atlas (ccv3). We first performed retinotopic mapping to define visual areas, and used the obtained maps to find, for each mouse, the optimal affine transformation to the Allen framework.

### Data analysis

All analyses of the behavioral effects of cortical inactivations were performed in Python 3.7. Generalized linear model (GLM) fitting of widefield data was performed in Matlab, and the results were analyzed in Python.

### Behavioral data selection

Because of the warm-up and easy-block trials, the sessions are naturally organized into a block structure, such that the duration of each block of the accumulating-towers task is of at least 40 trials (see above). We selected all trials from blocks in which the control (laser off) performance was at least 60% correct, collapsed over all levels of sensory evidence. After block selection, we excluded trials in which the animals failed to reach the end of the maze, or in which the total traveled distance exceeded the nominal maze length by more than 10% (Pinto et al., 2018, 2019). Additionally, because we were interested in assessing the effects of inactivation on accumulation timescales, we excluded animals that failed to use evidence from all quarters of the cue region of the maze to make their decisions in control trials. To do this, we fitted the logistic regression model (see below) separately for each animal, bootstrapping by sampling trials with replacement 200 times. We then computed the significance at each y position bin as the fraction of trials in which the model coefficient was equal to or greater than zero. Mice with any coefficients not significantly different than zero after false discovery rate correction (see below) were excluded from further analyses. These selection criteria yielded a total of 855 optogenetic inactivation sessions from 20 mice (average ∼43/mouse), corresponding to 100,787 control (laser off) trials, and 27,606 inactivation trials (average ∼511/condition, see Figure 1–table supplement 1). Twenty-five sessions from six mice were selected for widefield imaging data analysis.

### Analysis of behavioral data

#### Overall performance

We calculated overall performance as the percentage of trials in which the mice turned to side with the highest tower counts, separately for control and inactivation trials.

#### Running speed

Speed was calculated for each inactivation segment using the total x-y displacement. We compared laser-induced changes in speed to control trials from the same maze segment.

#### Psychometric curves

We computed psychometric curves separately for control and inactivation trials by plotting the percentage of right-choice trials as a function of the difference in the number of right and left towers (#R – #L, or Δ). Δ was binned in increments of 5 between -15 and 15, and its value defined as the average Δ weighted by the number of trials. We fitted the psychometric curves using a 4-parameter sigmoid:

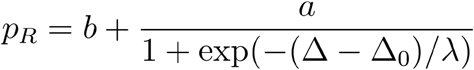

#### Evidence-weighting curves

To assess how mice weighted sensory evidence from different segments of the cue region, we performed a logistic regression analysis in which the probability of a right choice was predicted from a logistic function of the weighted sum of the net amount of sensory evidence from each of 4 equally-spaced segments (10 – 200 cm, since no towers can occur before y = 10):

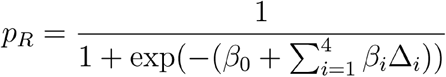

where Δ = # right – # left towers calculated separately for each segment. These weighting functions were calculated separately for ‘laser on’ and ‘laser off’ trials. To quantify the laser-induced changes in evidence weighting, we simply subtracted the ‘laser on’ from the ‘laser off’ curves, such that negative values indicate smaller evidence weights in the ‘laser on’ condition. Bin sizes were chosen to match the resolution of our inactivation epochs.

#### Laser-triggered analysis

For each area, we aligned the evidence-weighting curves by the position of laser onset, which was defined as y = 0, and used y position bins going up to 100 cm in the past. Thus, each inactivation condition contributed up to 3 bins of data, depending on the position of laser onset. For inactivations lasting more than one position bin (i.e. 100 cm), we used only the first bin during the inactivation as the y = 0 datapoint. Given the laser onset positions in our experiments, y = 0 had six data points, y = -50 had four, and y = -100 had three. For the laser-offset triggered analysis (Figure 2–figure supplement 2), we aligned the evidence-weighting curves by the first bin following laser offset. Given the laser offset positions in our data, we analyzed y = 50 (n = 4 conditions) and y = 100 (n = 3).

#### Statistics of inactivation effects

Error estimates and statistics for general performance, running speed and logistic regression weights were generated by bootstrapping this procedure 10,000 times, where in each iteration we sampled trials with replacement. P-values were calculated as the fraction of bootstrapping iterations in which the control-subtracted inactivation value was above zero. In other words, we performed a one-sided test of the hypothesis that inactivation decreases performance, speed and evidence weights on decision. To analyze the laser-onset triggered curves (Figure 2), we used a one-way ANOVA with repeated measures with the y position bin as a factor to establish the significance of the difference in the effects across bins. To account for the different number of datapoints per spatial bin, we implemented this as a mixed model with experimental conditions as the random effect. To assess whether laser effects were significant for each bin in the laser-onset (or offset)-aligned curves, we performed a one-sided t test against zero, with inactivation epochs as data points. Finally, to compare the effects of simultaneous and individual area inactivations (Figure 2–figure supplement 3), we performed a two-way ANOVA with repeated measures with factors y position bin and inactivation type, using the individual inactivation epochs as data points (in the case of individual inactivations, epochs from different areas were concatenated).

#### Clustering of evidence-weighting curves

We generated a 7 x 3 (areas x laser-onset-triggered inactivation bins) matrix containing the average laser-subtracted evidence-weighting curves, aligned by laser onset, for each individually targeted area. Thus, we excluded the experimental conditions in which frontal or posterior cortical areas were inactivated simultaneously from this analysis. We then performed spectral clustering into *k* clusters on that matrix. We tested *k* = 2 – 5, and chose the value of *k* that maximized clustering quality as measured by the Calinski-Harabasz index. Given the small number of areas per cluster, we generated error estimates for each y position bin by concatenating the individual inactivation conditions (epochs) for all areas of the cluster.

### Generalized linear model (GLM) of widefield data

We fitted Ca^2+^ activity averaged over each anatomically defined ROI with a generalized linear model (GLM)(Pinto et al., 2019; Pinto and Dan, 2015; Scott et al., 2017). For each trial and y position in the maze, we extracted ΔF/F (with native 10-Hz sampling frequency) limited to 0 ≤ y ≤ 300 cm (i.e. trial start, outcome and inter-trial periods were not included). Activity was then z-scored across all trials. ΔF/F of each area was modeled as a linear combination of different predictors at different time lags. In addition to the previously used task-event predictors (Pinto et al., 2019), we added coupling terms i.e. the zero-lag activity of the other simultaneously imaged ROIs (Pillow et al., 2008; Runyan et al., 2017), as well as auto-regressive terms to capture activity auto-correlations that were independent of task events (Spitmaan et al., 2020). Finally, we added a term to penalize the L2 norm of the coefficients, i.e. we performed ridge regression. The full model was thus defined as:

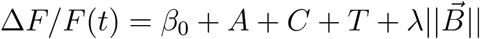

where *β*_*0*_ is an offset term, *λ* is the penalty term and 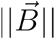 is the L2 norm of the weight vector. Additionally, *A, C* and *T* are the auto-regressive, coupling and task terms, respectively:

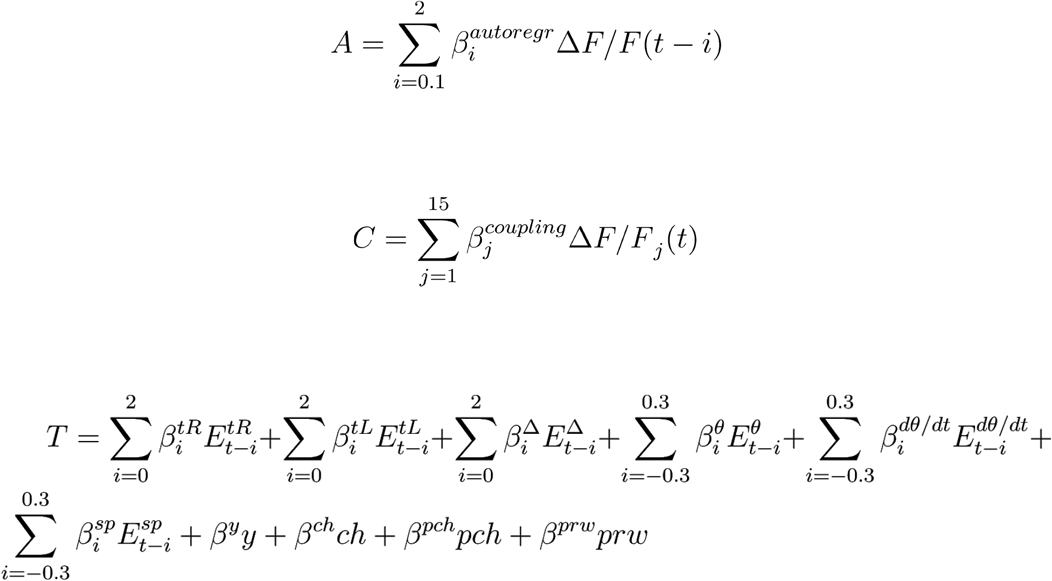

In the above equations, 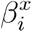 is the encoding weight for predictor *x* at time lag *i* (in steps of 0.1 s), where *x* is either a task event or the activity of the ROI at a previous time point, and 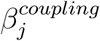 is the weight for the zero-lag activity for simultaneously imaged ROI *j* (we had a total of 16 ROIs across the two hemispheres). In the task term, 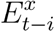 is a delta function indicating the occurrence of event *x* at time *t*-*i*. Specifically, *tR* indicates the occurrence of a right tower, *tL* of a left tower, *Δ* = cumulative #R – #L towers, *θ* is view angle, *dθ/dt* is virtual view angle velocity, *sp* is running speed, *y* is spatial position in the maze stem (no lags), and *ch, pch* and *prw* are constant offsets for a given trial, indicating upcoming choice, previous choice (+1 for right and –1 for left) and previous reward (1 for reward and –1 otherwise), respectively.

#### Cross-validation

The model was fitted using 3-fold cross-validation. For each of 20 values of the penalty term *λ*, we trained the model using ⅔ of the trials (both correct and wrong choices), and tested it on the remaining ⅓ of trials. We picked the value of *λ* that maximized accuracy, and used median accuracy and weight values across all 10 x 3 runs for that *λ*. Model accuracy was defined as the linear correlation coefficient between actual ΔF/F and that predicted by the model in the test set.

#### Model comparison

We tested three versions of the GLM, one with just the task term *T*, another one adding the auto-regressive term *A*, and the other with the coupling term *C* in addition to *A* and *T*. All versions were fitted using exactly the same cross-validation data partitioning to allow for direct comparison. We averaged cross-validated predictions over hemispheres and sessions for each mouse, performing the comparison with mouse-level data. Statistical significance of the differences between the accuracy of different models was computed using a two-way ANOVA with repeated measures with factors ROI and model type, and individual model comparisons were made using Tukey’s post-hoc test. Coefficient analysis in Figure 3 is from the full model, which had the highest performance.

### Quantification of timescales from the GLM

To quantify the timescales from the fitted auto-regressive coefficients, for each behavioral session we fitted an exponential decay function to the coefficients between 0.1 and 2 s in the past, normalized to the coefficient at 0.1 s (first bin):

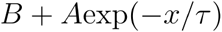

where *B* is the offset term, *A* controls the amplitude of the curve, *x* is the vector of normalized coefficients and *τ* is the decay time constant. Fits were performed using the non-linear least squares algorithm. The extracted time constants (τ) were first averaged over hemispheres and sessions for each mouse, and statistics were performed on mouse averages. Significance of the differences in the time constants across regions was assessed by performing a one-way ANOVA with repeated measures, with cortical regions as the factor.

### False discovery rate correction

We corrected for multiple comparisons using a previously described method for false discovery rate (FDR) correction (Benjamini and Hochberg, 1995; Guo et al., 2014; Pinto et al., 2019). Briefly, p-values were ranked in ascending order, and the *i*th ranked p*-*value, *P*_*i*_, was deemed significant if it satisfied *P*_*i*_ ≤ (*αi*)/*n*, where *n* is the number of comparisons and *α* is the significance level. In our case, *α* = 0.05 because we defined all tests as one sided.

## Data and code availability

Data analysis code and source code for figures is available at https://github.com/BrainCOGS/PintoEtAl2020_subtrial_inact.git. Behavioral data from inactivation experiments and GLM summary data will be deposited on a public repository upon peer-reviewed publication of this manuscript.

## Acknowledgements

We thank Sue Ann Koay and Kanaka Rajan for discussions, Abigail Russo and E. Mika Diamanti for comments on the manuscript, and Samantha Stein and Scott Baptista for technical assistance. This work was supported by the NIH grants U01NS090541, U19NS104648, F32NS101871 (L.P.) and K99MH120047 (L.P.).

## Author contributions

L.P. performed the experiments and analyzed the data; L.P. wrote the manuscript with input from C.D.B. and D.W.T.; L.P., C.D.B. and D.W.T. conceived the project.

## Table and Figure Supplements

**Figure 1–table supplement 1.** Numbers of mice, sessions and trials for each of the 54 experimental conditions. Last line shows the number of unique mice and trials across all experiments, as conditions were partially overlapping for a given mouse and behavioral session.

| Region | Epoch | Num. control trials | Num. laser trials | Num. mice | Num. sessions | Total trial count |
| --- | --- | --- | --- | --- | --- | --- |
| V1 | cue, 1<sup>st</sup> quarter (0 – 50 cm) | 21,741 | 692 | 7 | 189 | 22,433 |
| V1 | cue, 2<sup>nd</sup> quarter (50 – 100 cm) | 21,646 | 686 | 7 | 189 | 22,332 |
| V1 | cue, 1<sup>st</sup> half (0 – 100 cm) | 19,753 | 540 | 7 | 158 | 20,293 |
| V1 | cue, 3<sup>rd</sup> quarter (100 – 150 cm) | 21,580 | 680 | 7 | 185 | 22,260 |
| V1 | cue, 2<sup>nd</sup> half (100 – 200 cm) | 20,492 | 585 | 7 | 163 | 21,077 |
| V1 | delay (200 - 300 cm) | 20,013 | 596 | 8 | 162 | 20,609 |
| mV2 | cue, 1<sup>st</sup> quarter (0 – 50 cm) | 18,372 | 589 | 7 | 159 | 18,961 |
| mV2 | cue, 2<sup>nd</sup> quarter (50 – 100 cm) | 18,495 | 577 | 7 | 162 | 19,072 |
| mV2 | cue, 1<sup>st</sup> half (0 – 100 cm) | 14,446 | 367 | 3 | 121 | 14,813 |
| mV2 | cue, 3<sup>rd</sup> quarter (100 – 150 cm) | 18,429 | 594 | 7 | 160 | 19,023 |
| mV2 | cue, 2<sup>nd</sup> half (100 – 200 cm) | 14,449 | 353 | 3 | 119 | 14,802 |
| mV2 | delay (200 - 300 cm) | 15,058 | 366 | 5 | 124 | 15,424 |
| PPC | cue, 1<sup>st</sup> quarter (0 – 50 cm) | 15,929 | 403 | 6 | 133 | 16,332 |
| PPC | cue, 2<sup>nd</sup> quarter (50 – 100 cm) | 15,993 | 429 | 6 | 134 | 16,422 |
| PPC | cue, 1<sup>st</sup> half (0 – 100 cm) | 9,006 | 615 | 9 | 63 | 9,621 |
| PPC | cue, 3<sup>rd</sup> quarter (100 – 150 cm) | 15,950 | 419 | 6 | 134 | 16,369 |
| PPC | cue, 2<sup>nd</sup> half (100 – 200 cm) | 7,505 | 266 | 5 | 52 | 7,771 |
| PPC | delay (200 - 300 cm) | 8,947 | 616 | 6 | 67 | 9,563 |
| RSC | cue, 1<sup>st</sup> quarter (0 – 50 cm) | 15,923 | 422 | 6 | 134 | 16,345 |
| RSC | cue, 2<sup>nd</sup> quarter (50 – 100 cm) | 16,064 | 418 | 6 | 136 | 16,482 |
| RSC | cue, 1<sup>st</sup> half (0 – 100 cm) | 11,736 | 925 | 10 | 86 | 12,661 |
| RSC | cue, 3<sup>rd</sup> quarter (100 – 150 cm) | 15,952 | 432 | 6 | 134 | 16,384 |
| RSC | cue, 2<sup>nd</sup> half (100 – 200 cm) | 13,116 | 1,018 | 8 | 94 | 14,134 |
| RSC | delay (200 - 300 cm) | 9,533 | 729 | 8 | 70 | 10,262 |
| Posterior | cue, 1<sup>st</sup> quarter (0 – 50 cm) | 16,147 | 455 | 4 | 126 | 16,602 |
| Posterior | cue, 2<sup>nd</sup> quarter (50 – 100 cm) | 16,287 | 482 | 4 | 128 | 16,769 |
| Posterior | cue, 1<sup>st</sup> half (0 – 100 cm) | 14,439 | 355 | 3 | 119 | 14,794 |
| Posterior | cue, 3<sup>rd</sup> quarter (100 – 150 cm) | 16,252 | 474 | 4 | 127 | 16,726 |
| Posterior | cue, 2<sup>nd</sup> half (100 – 200 cm) | 14,038 | 312 | 3 | 118 | 14,350 |
| Posterior | delay (200 - 300 cm) | 14,117 | 357 | 3 | 115 | 14,474 |
| mM2 | cue, 1<sup>st</sup> quarter (0 – 50 cm) | 21,787 | 703 | 7 | 191 | 22,490 |
| mM2 | cue, 2<sup>nd</sup> quarter (50 – 100 cm) | 21,615 | 692 | 7 | 187 | 22,307 |
| mM2 | cue, 1<sup>st</sup> half (0 – 100 cm) | 19,972 | 588 | 7 | 161 | 20,560 |
| mM2 | cue, 3<sup>rd</sup> quarter (100 – 150 cm) | 21,766 | 674 | 7 | 190 | 22,440 |
| mM2 | cue, 2<sup>nd</sup> half (100 – 200 cm) | 21,015 | 634 | 8 | 167 | 21,649 |
| mM2 | delay (200 - 300 cm) | 19,755 | 605 | 7 | 159 | 20,360 |
| aM2 | cue, 1<sup>st</sup> quarter (0 – 50 cm) | 16,025 | 426 | 6 | 135 | 16,451 |
| aM2 | cue, 2<sup>nd</sup> quarter (50 – 100 cm) | 16,023 | 420 | 6 | 135 | 16,443 |
| aM2 | cue, 1<sup>st</sup> half (0 – 100 cm) | 20,338 | 576 | 7 | 166 | 20,914 |
| aM2 | cue, 3<sup>rd</sup> quarter (100 – 150 cm) | 16,046 | 425 | 6 | 135 | 16,471 |
| aM2 | cue, 2<sup>nd</sup> half (100 – 200 cm) | 21,210 | 658 | 8 | 169 | 21,868 |
| aM2 | delay (200 - 300 cm) | 19,737 | 616 | 7 | 158 | 20,353 |
| M1 | cue, 1<sup>st</sup> quarter (0 – 50 cm) | 13,894 | 369 | 6 | 117 | 14,263 |
| M1 | cue, 2<sup>nd</sup> quarter (50 – 100 cm) | 13,938 | 382 | 6 | 119 | 14,320 |
| M1 | cue, 1<sup>st</sup> half (0 – 100 cm) | 14,806 | 387 | 3 | 125 | 15,193 |
| M1 | cue, 3<sup>rd</sup> quarter (100 – 150 cm) | 14,009 | 364 | 6 | 121 | 14,373 |
| M1 | cue, 2<sup>nd</sup> half (100 – 200 cm) | 14,774 | 395 | 3 | 122 | 15,169 |
| M1 | delay (200 - 300 cm) | 14,721 | 386 | 3 | 123 | 15,107 |
| Frontal | cue, 1<sup>st</sup> quarter (0 – 50 cm) | 16,255 | 475 | 4 | 126 | 16,730 |
| Frontal | cue, 2<sup>nd</sup> quarter (50 – 100 cm) | 16,178 | 485 | 4 | 128 | 16,663 |
| Frontal | cue, 1<sup>st</sup> half (0 – 100 cm) | 14,410 | 369 | 3 | 120 | 14,779 |
| Frontal | cue, 3<sup>rd</sup> quarter (100 – 150 cm) | 16,065 | 483 | 4 | 123 | 16,548 |
| Frontal | cue, 2<sup>nd</sup> half (100 – 200 cm) | 14,392 | 377 | 3 | 121 | 14,769 |
| Frontal | delay (200 - 300 cm) | 14,511 | 365 | 3 | 118 | 14,876 |
| **Total unique count** |  | 100,787 | 27,606 | 20 | 855 | 128,393 |

**Figure 1–figure supplement 1.**
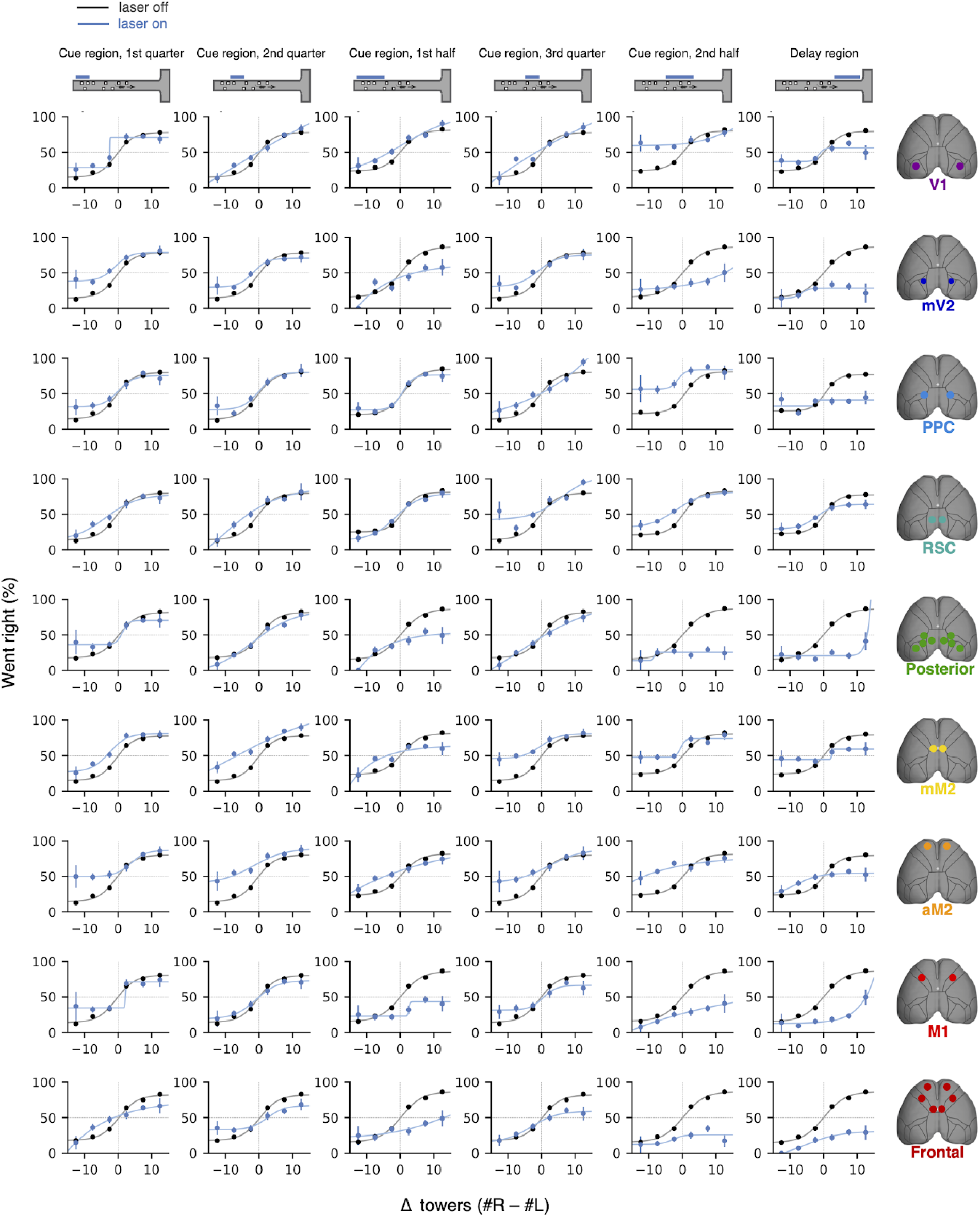
Effects of subtrial inactivations on psychometric functions during all 54 area-epoch combinations. Black lines show control trials and blue lines show inactivation trials, for data combined across mice. Error bars, binomial confidence intervals. Lines are best-fitting psychometric functions.

**Figure 1–figure supplement 2.**
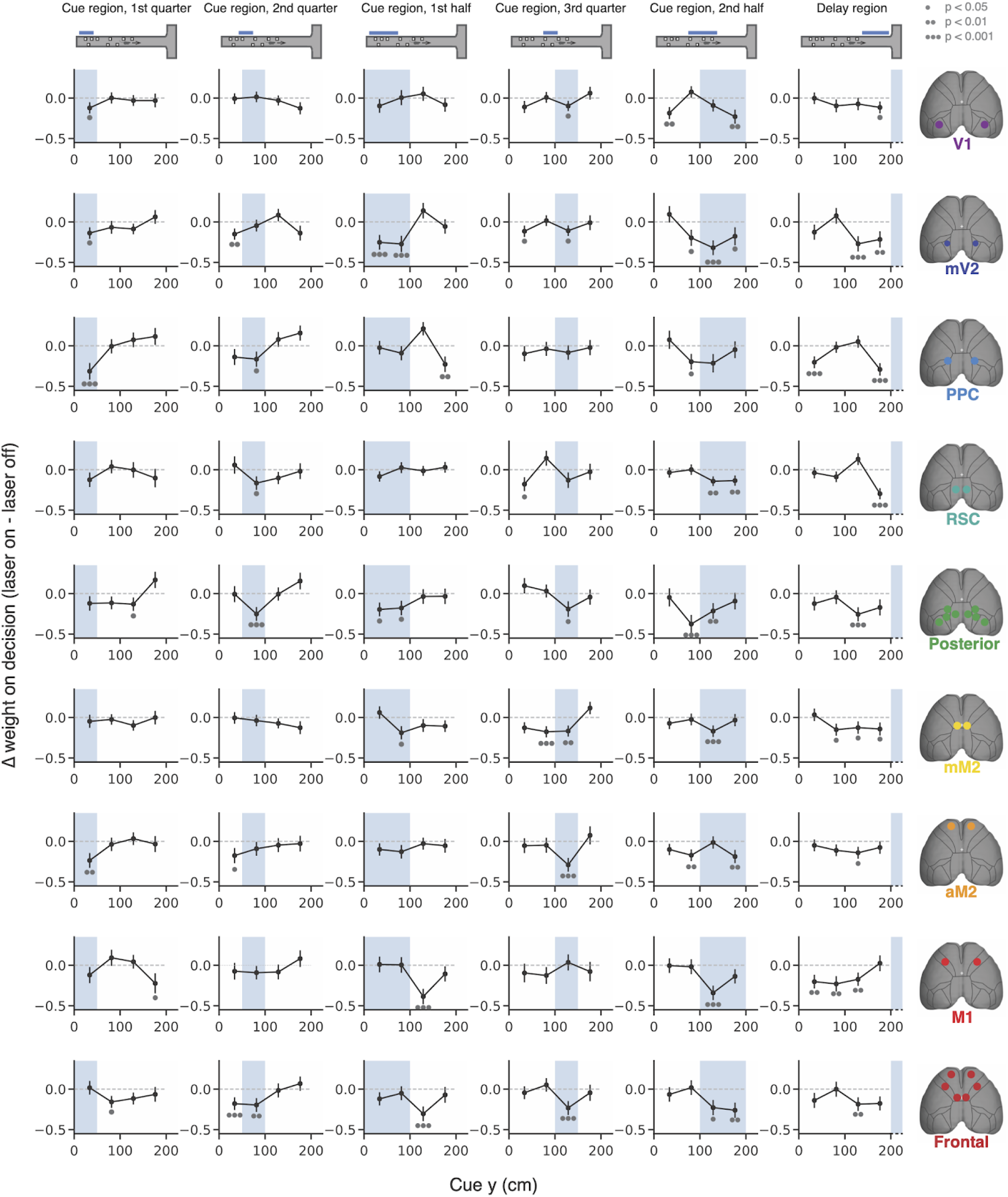
Effects of subtrial inactivations on evidence-weighting functions during all 54 area-epoch combinations. Black lines show the inactivation-induced change in evidence weights (laser on – laser off), and shaded areas indicate inactivation epoch. Data were combined across mice. Error bars, S.D. across 10,000 bootstrapping iterations. Gray circles indicate statistical significance according to the caption on top.

**Figure 2–figure supplement 1.**
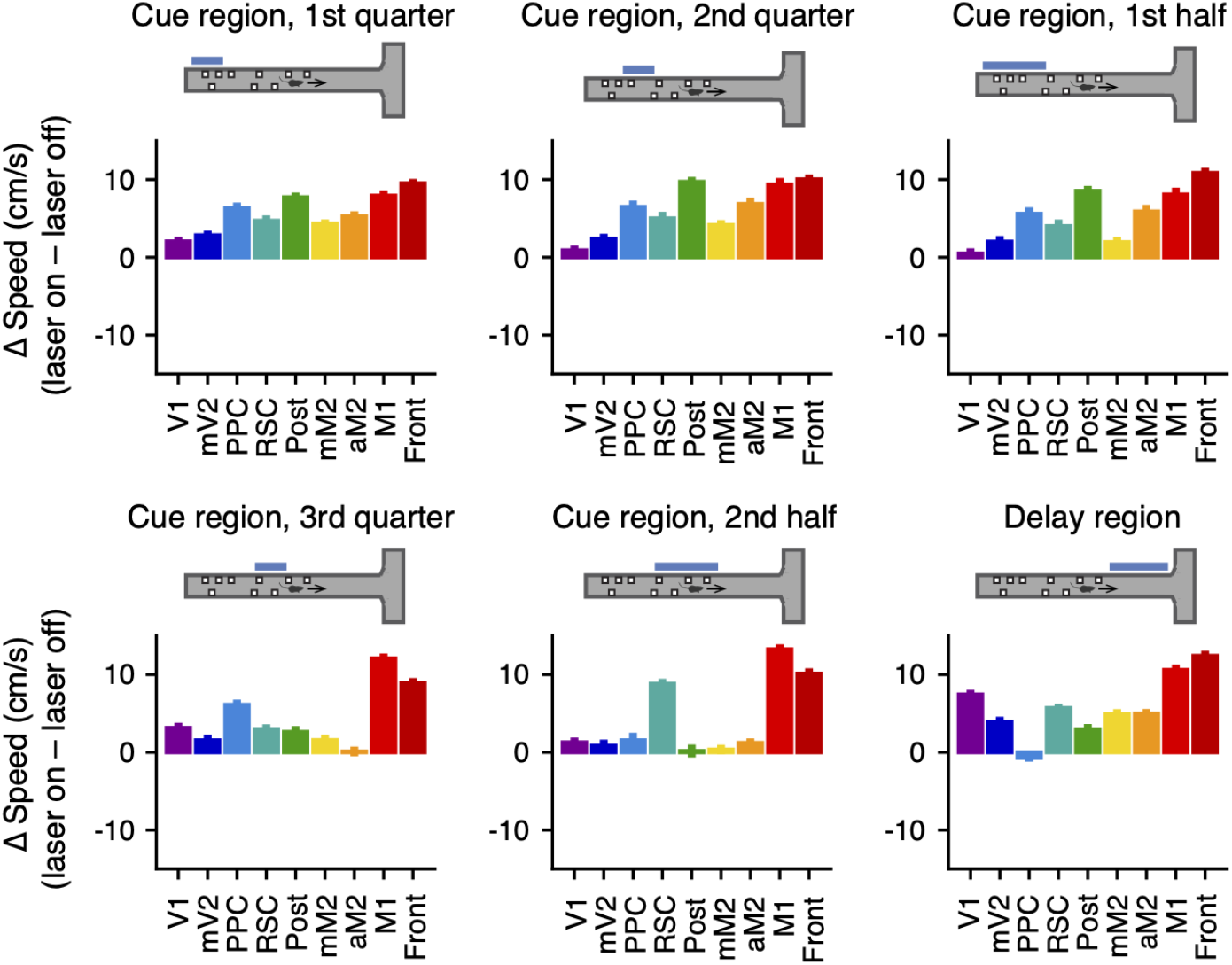
Inactivation of cortical areas does not decrease running speeds. Effects of subtrial inactivations on running speed during all 54 area-epoch combinations. Each panel shows inactivation-induced change in speed during the laser-on epoch or equivalent maze regions in control trials, for data combined across mice. Error bars: S.D. across 10,000 bootstrapping iterations. There were no significant decreases, as assessed by a one-sided bootstrapping test (see Materials and Methods).

**Figure 2–figure supplement 2.**
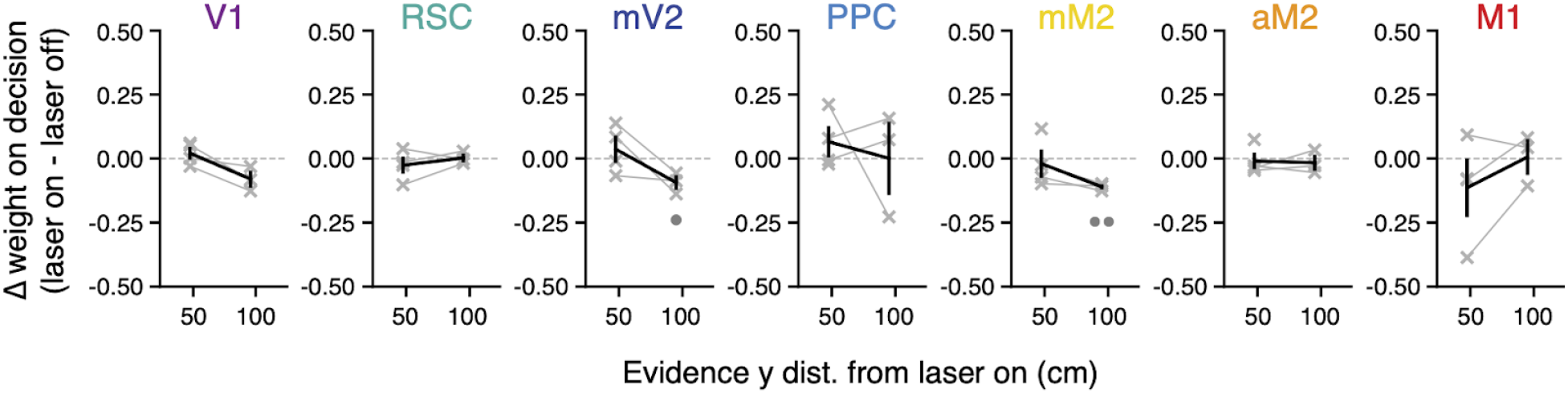
Little effect on evidence weighting after laser offset. Laser-offset-aligned changes in evidence-weighting curves for each bilaterally targeted area (laser on - laser off). Thin gray lines, individual inactivation epochs (n = 4 for y = 50, 3 for y = 100). Thick black lines, average across conditions. Error bars, ± SEM across experimental conditions, for data combined across mice. Circles below the lines indicate statistical significance (one circle: p < 0.05, two circles: p < 0.01; one-sided paired t test vs. zero, corrected for multiple comparisons).

**Figure 2–figure supplement 3.**
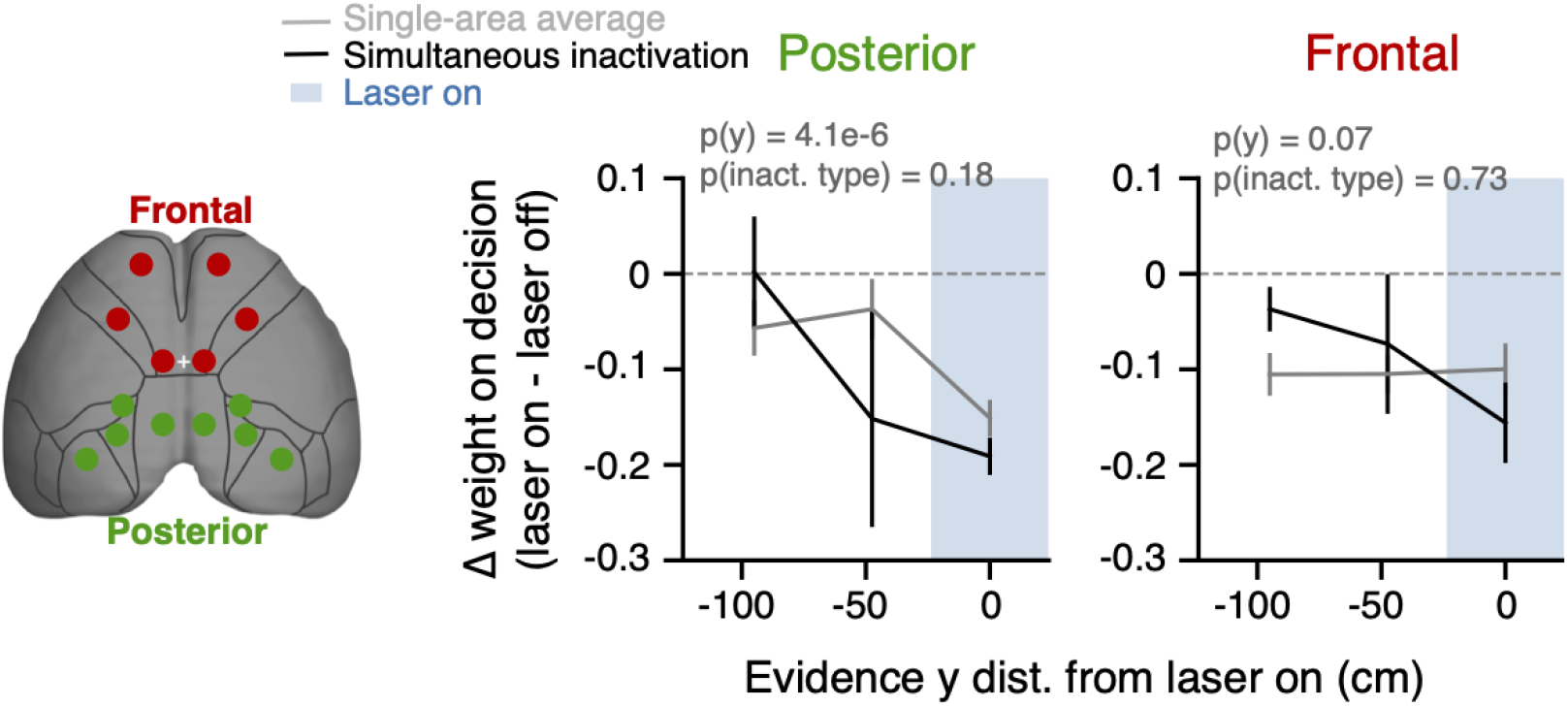
Comparison between simultaneous multi-area inactivation and average across the corresponding individually inactivated areas. Average laser-onset-aligned changes in evidence-weighting curves for each set of experimental conditions (caption on top), for data combined across mice. Error bars, ± SEM across experimental conditions. Shaded areas, laser on periods. P-values on top are from a two-way ANOVA with repeated measures, factors inactivation type (simultaneous vs. individual) and y position. There were no differences between simultaneous and individual inactivation of either frontal or posterior areas. On the other hand, effects depended on the distance of evidence from laser onset for posterior but not frontal areas.

**Figure 3–figure supplement 1.**
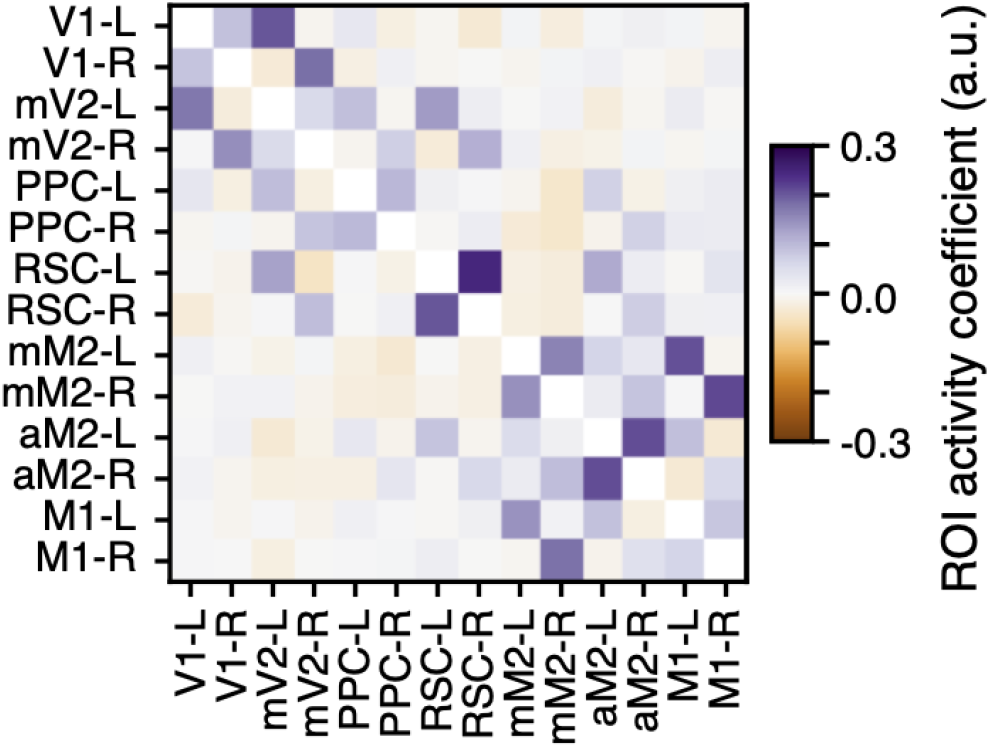
GLM coupling coefficients for ROI activity predictors. For each ROI (rows), shown are the average coefficients (n=6 mice) for the predictors consisting of zero-lag activity of the other simultaneously imaged ROIs. Note that the diagonal elements are not defined since zero-lag self-activity predictors were not in the model. Data from the somatosensory cortex ROI has been omitted for symmetry with the inactivation data. L and R indicate left and right hemispheres, respectively.

